# Additive and epistatic effects influence spectral tuning in molluscan retinochrome opsin

**DOI:** 10.1101/2021.05.26.445805

**Authors:** G. Dalton Smedley, Kyle E. McElroy, Jeanne M. Serb

## Abstract

The relationship between genotype and phenotype is nontrivial due to often complex molecular pathways that make it difficult to unambiguously relate phenotypes to specific genotypes. Photopigments, an opsin apoprotein bound to a light-absorbing chromophore, present an opportunity to directly relate the amino acid sequence to an absorbance peak phenotype (λ_max_). We examined this relationship by conducting a series of site-directed mutagenesis experiments of retinochrome, a non-visual opsin, from two closely related species: the common bay scallop, *Argopecten irradians*, and the king scallop, *Pecten maximus.* Using protein folding models, we identified three amino acid sites of likely functional importance and expressed mutated retinochrome proteins *in vitro.* Our results show that the mutation of amino acids lining the opsin binding pocket are responsible for fine spectral tuning, or small changes in the λ_max_ of these light sensitive proteins Most mutations caused a blue shift regardless of the retinochrome background, with shifts ranging from a 12 nm blue shift to a 5 nm red shift from the wild-type λ_max_. These mutations do not show an additive effect, but rather suggests the presence of epistatic interactions. This work highlights the importance of binding pocket shape in the evolution of spectral tuning and builds on our ability to relate genotypic changes to phenotypes in an emerging model for opsin functional analysis.

**Author summary:** Site-directed mutagenesis determined that spectral tuning in retinochrome is not solely additive, but is influenced by intra-molecular epistasis.

## Introduction

Understanding how genotype influences phenotype is a fundamental goal of biology. This relationship is far from trivial, as complex genetic pathways and environmental effects often preclude clear connections. The phototransduction system that converts light into electrical signals [1] offers a rare opportunity to examine genotype-phenotype relationships because single amino acid substitutions can drastically alter visual protein phenotype [2]. Furthermore, because photopigments – the functional unit of phototransduction – only absorb a portion of the light spectrum, the maximum absorbance (λ_max_) represents a directly measurable phenotype.

Photopigments form by covalent bonds between a retinal chromophore and the binding pocket of an opsin apoprotein. The interactions of the chromophore with amino acids in the binding pocket have been highlighted as critical for photopigment function by theoretical calculations [3,4], mutagenesis [5–9] and quantum mechanics/molecular mechanics (QM/MM) modeling [10–12]. Such interactions between the chromophore and the amino acid residues within the binding pocket of the opsin are thought to be responsible for modulating λ_max_, or spectral tuning [3,13]. Changing amino acid side chains that effect the environment of the chromophore binding pocket may be particularly important [14].

Extensive mutagenesis efforts have been directed towards identifying spectral tuning sites in opsins [15–18]. In vertebrates, λ_max_ differences between middle- and long-wavelength-sensitive pigments involved in color vision are attributed to three [19–21] or five [22,23] amino acid sites. These individual sites are responsible for small nanometer shifts that cumulatively alter the protein function [7]. Whether the patterns of amino acid changes responsible for spectral tuning in vertebrate visual opsins are broadly applicable to the immense diversity of opsin proteins is unknown. This is because the vast majority of experimental spectral tuning research has been done in vertebrate systems (e.g., monostable G_t_-protein coupled opsins) (reviewed in [24,25]), with few exceptions in non-vertebrate species (e.g., bistable G_q_-protein coupled opsins [26–28]). Moreover, the crystal structure of invertebrate opsins, such as G_q_-protein coupled opsins (e.g., squid “rhodopsin”), have important differences compared to vertebrate opsin (G_t_-protein coupled opsins, e.g., bovine rhodopsin), such as greater organization in the cytoplasmic region correlating to bistability, an interconvertibility between the photopigment’s dark state and its photoproduct [29,30]. This longstanding focus on vertebrate opsins has left us with a severe lack of understanding for how spectral tuning functions in invertebrate opsins.

To identify amino acids involved in the spectral tuning of photopigments, comparing the amino acid sequence and protein function of two closely related species is an effective starting point. Here, we cloned and expressed *in vitro* a mollusc-specific opsin, retinochrome, from two closely related scallops: the common bay scallop, *Argopecten irradians* and the king scallop, *Pecten maximus.* Retinochrome is one of the few examples of a non-vertebrate, non-visual opsin with well-characterized biochemical and spectral properties [31–34]. Further, its ease of expression and predictable nature in heterologous cell culture makes it a compelling protein to work with [31,35–37]. Retinochrome belongs to the “photoisomerase” opsin clade of RGR/retinochrome/peropsin [38], which binds to the all-*trans* isomer of retinal as opposed to 11-*cis* isomer commonly used by most visual opsins. The retinochrome counterion that stabilizes chromophore binding has a notably different location than other opsins [39]. Furthermore, the likely role of retinochrome is as a photoisomerase in the retinoid visual cycle of molluscs, converting all-*trans* retinal to 11-*cis* retinal following the absorption of light [40], which sets it apart from opsins that mediate phototransduction. Whether a few amino acid substitutions cumulatively contribute to differences in retinochrome λ_max_ values, as seen in vertebrate visual opsins, is unknown.

We hypothesize that changing the shape [41] or electrostatic environment [42] of retinochrome’s binding pocket will shift the λ_max_ value. This shift would be the result of different biochemical properties of newly introduced amino acid residues, which can affect noncovalent interactions within the photopigment. To test this hypothesis, we performed reciprocal site-directed mutagenesis of retinochromes from the common bay scallop, *Argopecten irradians*, and the king scallop, *Pecten maximus*. Using a protein modeling approach, we identified three amino acid sites in the binding pocket and created 14 mutants representing all possible combinations of the three sites of interest in each background. Spectral analyses indicate that these three sites have differing degrees of influence on phenotype and appear to have epistatic interactions with other regions of the protein. Our results reveal a more complicated path from genotype to phenotype than typically seen for opsins.

## Results

### Retinochromes have similar sequence, but differ spectrally, between scallop species

Cloned retinochrome sequence of *Argopecten irradians* (GenBank KX550908) and *Pecten maximus* (GenBank MW984523) were 308 amino acids. The two protein sequences were 92% similar, with 25 different amino acid residues. There was high sequence conservation within the binding pocket (99%), and most of the residue differences belonged to the same biochemical class. Residues surrounding sites important for the function of retinochrome, such as the lysine for chromophore binding (pos 274) and glutamate counterion (pos 160), are conserved between scallop samples, as are amino acids around these sites.

To determine the λ_max_ of *A. irradians* retinochrome (Airr-RTC) and *P. maximus* retinochrome (Pmax-RTC), we expressed both proteins in HEK293T cells and incubated with all-*trans* retinal. The absorbance peak (λ_max_) of Airr-RTC was 510 nm (Fig 1a, black curve), while the absorbance peak of Pmax-RTC was 492 nm (Fig 1b, black curve). This is the first *in vitro* measurement of retinochrome λ_max_, and our measurements are consistent with prior literature [33,43]. Differences between the dark spectra minus spectral tracings of the photopigment after exposure to blue light (Fig 1a-b, red curves) confirm the formation of a photopigment (Fig 1, insets).

**Fig 1.**
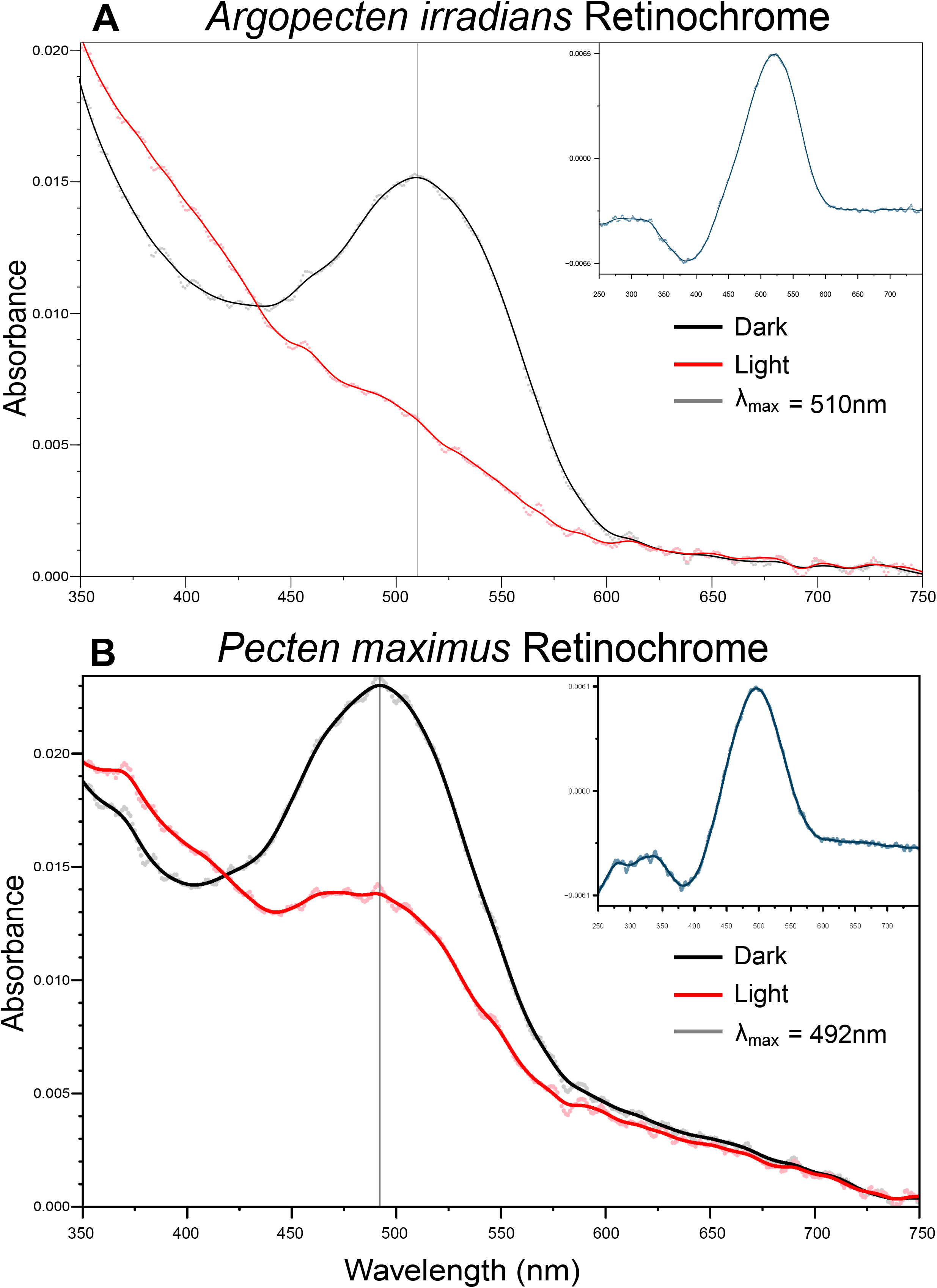
Absorbance spectra of Airr-RTC (A) and Pmax-RTC (B) after *in vitro* expression and purification. Dark spectra (black curves) represent the average absorbance from five spectral measurements (dots) of the photopigment prior to its exposure to light. Grey vertical lines of each dark spectrum indicate the λ_max_. Red curves are the average absorbance from five spectral measurements (red dots) of the photopigment after 3-minute exposure to blue light. The inset shows the differential absorbance of the dark spectrum minus the spectrum recorded after irradiation with blue light.

### Three amino acid changes indicate potential interaction sites lining the retinal binding pocket

To narrow down the candidate list of 25 spectral tuning sites, we employed three-dimensional protein models to identify potential ligand interaction sites. In total, 18 positions were identified between the two retinochrome models and were at the same binding pocket locations of two RTC homologs with two exceptions. Positions 170 and 188 were only predicted to be interaction sites in the Pmax-RTC model, but not in the Airr-RTC model (Fig 2a, 2b). However, the amino acid residue was conserved at position 170 and thus was removed from the candidate list. In contrast, position 188 varied between a tryptophan in Pmax-RTC versus a tyrosine in Airr-RTC. While both of these amino acids are classified as hydrophobic with similar pKa scores, the position of the tyrosine’s hydroxyl group could allow a more direct polarity change in the binding pocket [44], and therefore may affect the absorbance of the photon by the bound retinal (Fig 2c). We selected site 188 as the first site for mutation.

**Fig 2.**
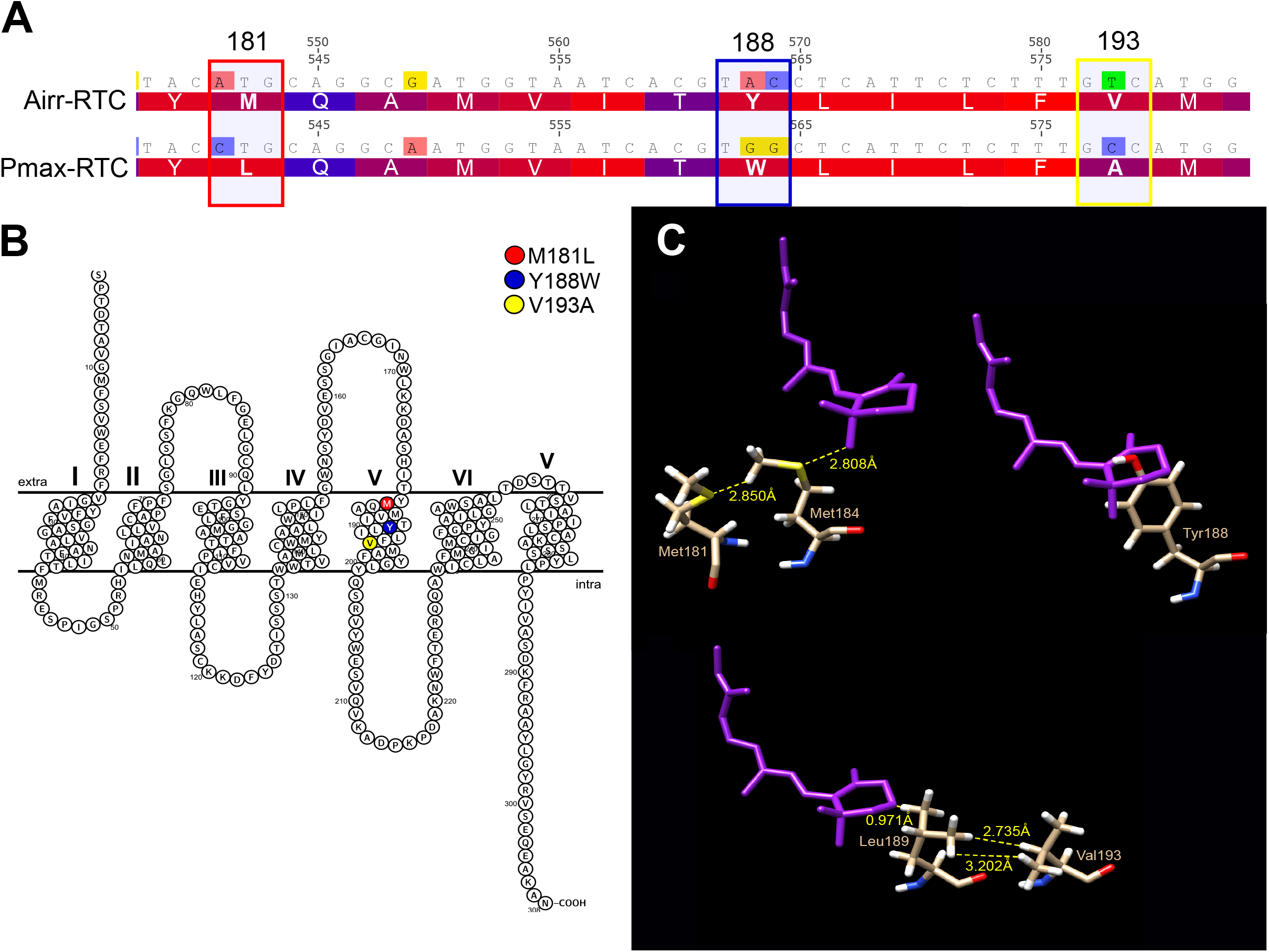
Location of candidate spectral tuning sites in retinochrome. (A) Amino acid and nucleotide sequence alignments of Airr-RTC and Pmax-RTC with nucleotide substitutions shaded. Large numbers are amino acid sites; small numbers are nucleotide positions of the coding sequence. Amino acids are colored by the degree of hydrophobicity of the side chains ranging from red (more hydrophobic) to blue (less hydrophobic). Colored boxes highlight the 3 amino acid differences between Airr-RTC and Pmax-RTC at sites with predicted involvement in spectral tuning and target nucleotide positions for mutagenesis. (B) Snakeplot showing secondary structure of Airr-RTC. The sites of the mutagenesis experiments are highlighted in red (M181), blue (Y188), and yellow (V193). Seven transmembrane helices are indicated by roman numerals. (C) 3D modeling of bound retinal chromophore and sites of interest. The 11-*cis* retinal molecule is purple, with the amino acid chains colored by atom composition. Yellow dash lines and numbers show the distances between structures in angstroms. Amino acid residue identities and location are labeled in beige next to the residue.

To identify other sites that may be responsible for altering the shape of or electrostatic environment within the binding pocket, we compared the 3D models of the Airr-RTC and Pmax-RTC focusing on the location of the non-conserved amino acids. We used UCSF Chimera to manipulate the 3D models, allowing us to measure the distance between active groups of the amino acid residues of interest in relation to one another as well as the retinal molecule. Many non-covalent polar interactions range from 0.5 to 3.5 angstroms, thus this range was used as a criterion for identifying sites of interest through bond measurements. Site 184 (Figure 2a, b) is a conserved methionine in both Pmax-RTC and Airr-RTC but was predicted by COACH to function as a ligand interaction site. A neighboring site which is not conserved is site 181. Site 181 is a methionine in Airr-RTC and a leucine in Pmax-RTC. The primary difference between these nonpolar side chains is the presence of the thioether of the methionine. This thioether allows for oxidation of the residue and acts as a strong hydrogen bond acceptor. Bond measurements show that the sulfur of Met181 is only 2.85 angstroms from the side chain hydrogens of Met184 (Figure 2c). We selected site 181 as a second site of interest due its potential in changing the polarity of an amino acid lining the binding pocket, site 184.

Using the same logic, we also identified nonconserved position 193 as a site of interest. Position 189, a leucine, was predicted by COACH as an interaction site for the ligand (Figure 2a, b), yet was conserved between the RTC homologs. However, within 0.5 to 3.5 angstroms, position 193 differed as a valine in Airr-RTC or an alanine in Pmax-RTC. The isopropyl group of the valine is much bulkier than that of the methyl group found in alanine. Given the proximity of the hydrogens found on the carbon groups of the leucine to that of the valine (between 2.735 angstroms and 3.202 angstroms) it is likely that the location of predicted interaction site 193 is altered in the space of the protein (Figure 2c).

### Selected amino acid mutations cause shifts in the maximum absorbance (λ_max_)

Single mutations had specific effects on λ_max_ in comparison to the Airr-RTC and Pmax-RTC wild-type proteins (Table 1). In Airr-RTC single mutants, site 181 (M181L) absorbed at 512 nm, a 2 nm change from the wild-type (Figure 3), while site 188 (Y188W) and site 193 (V193A) resulted in blue shift of either 5 or 12 nm (λ_maxes_ of 505 nm and 498 nm), respectively (Figure 3). The blue shift from mutations at 188 and 193 suggest their role as spectral tuning sites. Both double mutants containing mutations in site 188 (Airr-RTC-ΔM181L-ΔY188W and Airr-RTC-ΔY188W-ΔV193A) also shifted blue, with λ_maxes_ of 498 nm and 502 nm, respectively (Figure 3). The double mutations at sites 181 and 193 had little change in the absorbance peak from the wild-type with a λ_max_ of 512 nm (Figure 3) in the Airr-RTC background. Considering the λ_max_ of the single mutants at sites 181 and 193, the combination of the two shows the effects at site 181 either compensate for or outweigh the effects at site 193. The triple mutant (Airr-RTC-ΔM181L-ΔY188W-ΔV193A) resulted in a 10 nm blue-shift (λ_max_ = 500nm) (Figure 3).

**Table 1.**
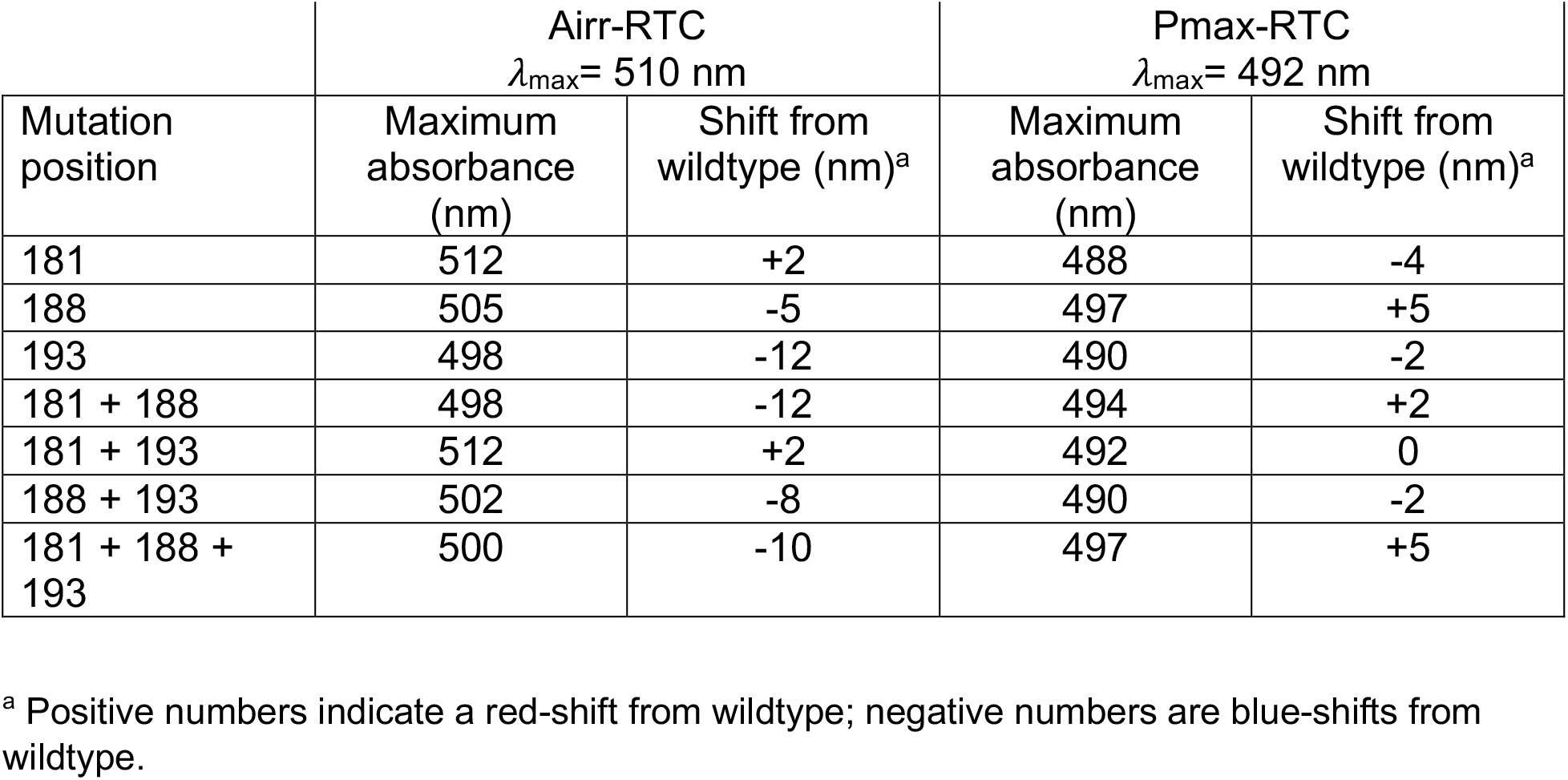
Mutant retinochrome **λ**_**max**_ values and the change in nanometers between mutant and wildtype **λ**_**max**_ values.

| Mutation position | Airr-RTC<br>$\lambda_{\text{max}} = 510 \text{ nm}$ | | Pmax-RTC<br>$\lambda_{\text{max}} = 492 \text{ nm}$ | |
| --- | --- | --- | --- | --- |
|  | Maximum absorbance (nm) | Shift from wildtype (nm)<sup>a</sup> | Maximum absorbance (nm) | Shift from wildtype (nm)<sup>a</sup> |
| 181 | 512 | +2 | 488 | -4 |
| 188 | 505 | -5 | 497 | +5 |
| 193 | 498 | -12 | 490 | -2 |
| 181 + 188 | 498 | -12 | 494 | +2 |
| 181 + 193 | 512 | +2 | 492 | 0 |
| 188 + 193 | 502 | -8 | 490 | -2 |
| 181 + 188 + 193 | 500 | -10 | 497 | +5 |

<sup>a</sup> Positive numbers indicate a red-shift from wildtype; negative numbers are blue-shifts from wildtype.

**Fig 3.**
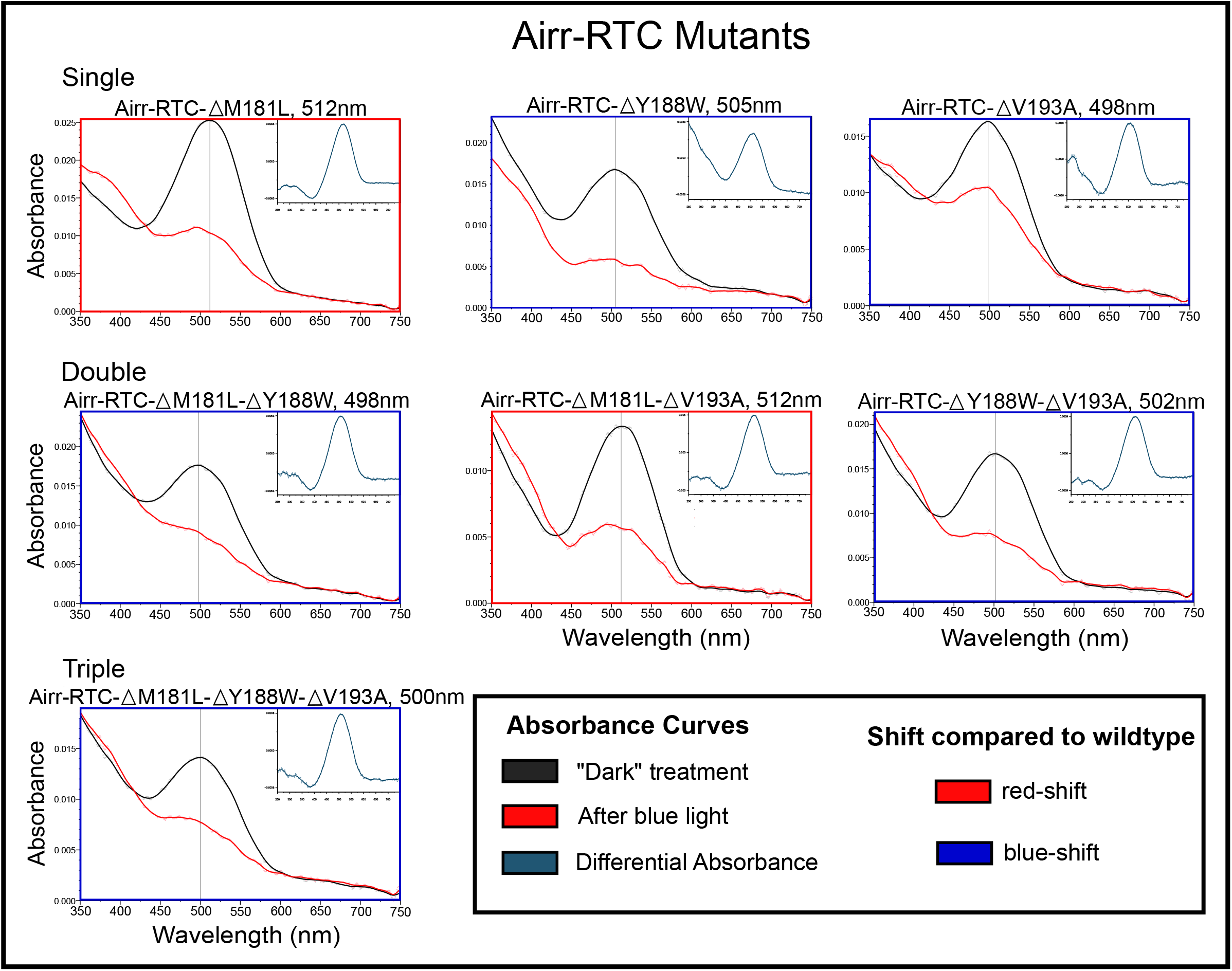
Site-directed mutagenesis cause spectral shifts in *Argopecten irradians* retinochrome. Black curves show the plot of dark (unexposed) spectra and red curves show absorption after 3-minute exposure to blue light. Vertical black lines highlight the maximum absorption peaks of mutants, the names of which reside above the spectra along with the value of the λ_max_ in nanometers (nm). Colored boxes around the spectra show the shift (blue or red) compared to the wild-type Airr-RTC. Insets show the differential absorbance of the dark spectrum minus the spectrum recorded after irradiation with blue light.

We also expressed and recorded the spectra of the seven mutant Pmax-RTC (Figure 4). From the single mutants, sites 181 (Leu to Met) and 193 (Ala to Val) showed a slight blue shift with λ_maxes_ of 488 nm and 490 nm, respectively (Figure 4). Site 188 (Trp to Tyr) showed a red shift to 497 nm(Figure 4). The Pmax-RTC double mutants had no or little change in λ_max_ from wild-type spectra. Double mutant protein Pmax-RTC-ΔM181L-ΔY188W had a λ_max_ of 494 nm, combination mutant Pmax-RTC- ΔM181L-ΔV193A showed no change in λ_max_ from the wild-type protein at 492 nm, and Pmax-RTC- ΔY188W-ΔV193A had a λ_max_ of 490 nm (Figure 4). The triple mutant Pmax-RTC-ΔM181L-ΔY188W-ΔV193A also showed a 5 nm red shift (λ_max_ = 497 nm; Figure 4).

**Fig 4.**
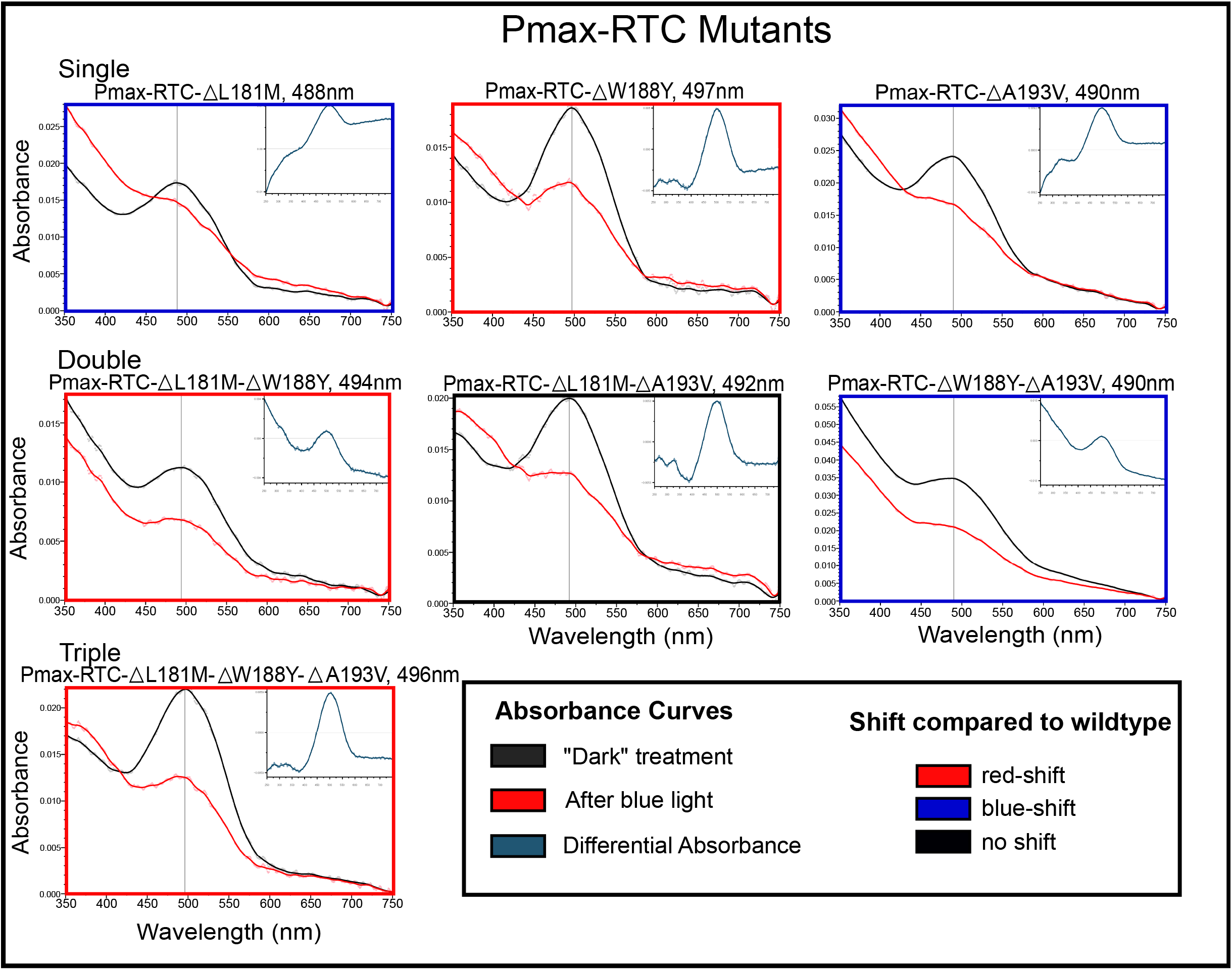
Site-directed mutagenesis cause spectral shifts in *Pecten maximus* retinochrome. Black curves show the plot of dark (unexposed) spectra and red curves show absorption after 3-minute exposure to blue light. Vertical black lines highlight the maximum absorption peaks of mutants, the names of which reside above the spectra along with the value of the λ_max_ in nanometers (nm). Colored boxes around the spectra show the shift (blue or red) compared to the wild-type Pmax-RTC with a black box indicating no change in the λ_max_ from wildtype. Insets show the differential absorbance of the dark spectrum minus the spectrum recorded after irradiation with blue light.

## Discussion

The relationship between genotype and phenotype is a central unanswered question in biology. Here, we provide the first experimental results with respect to absorbance spectra of a molluscan photopigment. We investigated the genotype-phenotype relationship by comparing retinochrome from two closely related scallop species and, through site-directed mutagenesis experiments, altered amino acids predicted to influence spectral absorbance. Because the *A. irradians* and *P. maximus* retinochrome sequences are highly conserved around both the Schiff base binding site and the counterion, the 18 nm difference in retinochrome λ_max_ that we observed between the two species must be attributed to non-conserved sites elsewhere in the protein. We hypothesized that amino acids lining the binding pocket may in part be responsible for spectral tuning of retinochrome λ_max_. We used bioinformatic methods to identify three non-conserved amino acids that may alter the shape or electrostatic environment of the opsin binding pocket, and reciprocally mutated those sites between *A. irradians* and *P. maximus* to test their role in spectral tuning. These three sites are, in part, responsible for spectral tuning of scallop retinochromes. However, our results show that the relationship between amino acid changes and spectral shifts is not simply additive, and indicates a role of intramolecular epistasis in retinochrome spectral tuning.

### Predicted Ligand Interaction Sites Alter λ_max_

Through protein modeling and subsequent testing of 14 reciprocal site-directed mutants, we found three sites that effect spectral tuning of retinochrome. However, these results were not always predictable or additive in nature. Site 188, a tyrosine in *A. irradians* and a tryptophan in *P. maximus*, was the only non-conserved predicted interaction site based on the COACH analysis. Mutations created at this site resulted in a -5 nm blue shift and a +5 nm red shift of λ_maxes_ for Airr-RTC-ΔY188W and Pmax-RTC-ΔW188Y, respectively. This mirroring effect between reciprocal mutants strongly indicates an independent effect on spectral tuning at this location. The impact of hydroxyl-containing amino acids, such as tyrosine, on λ_max_ is well documented in both vertebrate and invertebrate photopigments [15,44,45]. Such studies demonstrate that the change in λ_max_ is due to the potential dipole moments created by the oxygen of hydroxyl groups in proximity of the *ß*-ionone ring of the chromophore, which cause a red shift of the absorption maximum or dipole residues around the protonated Schiff base that result a λ_max_ shift in the opposite direction (e.g., OH-site rule [46]). The former may be occurring in Pmax-RTC-ΔW188Y. Our Airr-RTC 3D model shows the tyrosine hydroxyl group inserted into the aromatic ring of the retinal, while the tryptophan in the Pmax-RTC model does not extend as far. This impossible orientation of tyrosine is likely the result of the software’s limitation, which only provides 11-*cis* retinal as a ligand option, rather than the all-*trans* retinal preferred by retinochrome. Likely, the straight poly-carbon chain of the all-*trans* retinal would rest differently in the binding pocket of the retinochrome, but the potential for dipole moments from the hydroxyl group could still affect the electrostatic environment of the binding pocket. This hydroxyl group is absent in the tryptophan at site 188 of wildtype *P. maximus*. Spectral analyses of the single, double, and triple mutants of Airr-RTC with the Y188W mutation all had a blue shift in λ_max_ ranging from -7 nm to -14 nm as expected under the OH-site rule, while spectra of Pmax-RTC single, double, and triple mutants with the W188Y mutation did not show the reciprocal pattern. The Pmax-RTC W188Y single mutant, L181M and W188Y double mutant, and L181M, W188Y and A193V triple mutant had slight red shifts between +2 nm to +5 nm, while the double mutant containing W188Y and A193V showed a blue shift of 2 nm. These conflicting changes in λ_max_ indicate that other polar groups may be involved in retinochrome spectral tuning. Future work should consider the “dipole-orientation rule,” an extension of the OH-site rule, that takes into account both location and orientation of polar groups, including water molecules, in modeling approaches [47].

The remaining two mutant sites, 181 and 193, allowed us to investigate the effects of residues in close proximity to predicted interaction sites rather than sites directly interacting with the retinal chromophore. The results of our site-directed mutagenesis experiments point to a hierarchy of effects. For instance, we observed the same +2 nm (λ_max_ 512 nm) shift in Airr-RTC with the M181L single mutation as with the double mutant with M181L and V193A, yet the single mutant V193A had a large -12 nm blue shift (λ_max_ 498 nm). The lack of spectral shift in the double mutant between these sites indicate that the effect posed by M181L may compensate for the alteration caused by the mutation at V193A in Airr-RTC backbone. In Pmax-RTC mutants, single mutants at same sites L181M and A193V had minor blue shifts (−4 nm and -2 nm, respectively), while the double mutant of those sites showed no change in λ_max_ (492nm). However, this blue shift was maintained in Pmax-RTC mutants at the same sites. The mutation with hydrophobic residues in bacterial rhodopsins have been shown to cause dramatic blue shifts (up to 80 nm relative to wild type) [48], because the side chains of these hydrophobic residues tend to be bulky due to the saturation of hydrogens and may restrict movement of the retinal’s *ß*- ionone ring. Thus, the presence of hydrophobic residues may alter the binding pocket geometry by pushing adjacent residues, such as those lining the binding pocket, into different orientations.

### A possible role for binding pocket shape in determining λ_max_

The effect of mutation V193A in Airr-RTC may exhibit a stronger effect on the shape of the binding pocket. The single Airr-RTC mutants containing V193A showed a large blue shift (−12 nm), with the exception of the double mutant also containing M181L, which showed a slight +2 nm red shift. The decrease in size of side chains from valine to alanine may be significant enough to reduce the bulging effect of this amino acid residue on the proposed ligand binding site, creating variation in the region of the protein and the shape or size of the binding pocket. However, when combined with M181L mutation, the small change of λ_max_ could suggest that the second mutation compensates the effect of V193A. Reciprocal mutants at the site 193 in Pmax-RTC (A193V) were expected to result in red shifts, mirroring that of Airr-RTC mutants, but these mutants also had consistent and large blue shifts (−10 nm to -12 nm) with the exception of the triple mutant (−5 nm). Here, the increase in side chain bulk (A to V) producing the blue shift in λ_max_ may indicate a weaken the interaction between the ligand and retinochrome.

A scenario where mutations in the binding pocket weaken the interactions with the chromophore may arise from changes in the distance between the amino acids in the binding pocket and the bound retinal as well as the distance between interacting amino acids. A change in distance can alter the strength of the interactions of these components thus potentially changing the polar environment of the binding pocket and the λ_max_ of the protein. While mutations in Airr-RTC appear to shift towards the wild-type λ_max_ of Pmax-RTC, the λ_max_ of the reciprocal mutants in Pmax-RTC may actually reflect reduced interaction strength between binding pocket residues and chromophore, thus shifting towards the native λ_max_ of unbound all-*trans* retinal in the UV region. However, being able to visualize the protein and binding pocket shape via crystal structure would help us to understand how the shape is being affected by specific mutations.

### Epistatic effect on spectral tuning

In numerous vertebrate visual opsins, spectral tuning is attributed to one or few amino acid substitutions [17]. For example, three amino acids are responsible for red-shifting in the evolution of old world primate trichromatic vision *via* additive effects ([49,50] but see [19]). We therefore had reason to expect one or particular combinations of the three predicted sites for spectral tuning to be sufficient for shifting λ_max_ from one homolog to the to the other. However, our reciprocal site-directed mutagenesis rarely mirrored the spectral tuning between Airr-RTC and Pmax-RTC backgrounds. In fact, site 188 was the only mutant with equal blue and red shifts for Airr-RTC (−5 nm) and Pmax-RTC (+5 nm), respectively, relative to their wildtype proteins.

Both triple mutants had λ_max_ shifts in the expected direction, though neither shift had a magnitude great enough to match the wildtype of the other species, *e.g.* Airr-RTC-ΔM181L-ΔY188W-ΔV193A λ_max_ did not match the Pmax-RTC λ_max._ The Pmax-RTC-ΔL181M-ΔW188Y and Pmax-RTC-ΔW188Y-ΔA193V double mutants had λ_max_ shifts that were close (within 1 nm) to the sums of λ_max_ shifts for the single mutants. Otherwise, the spectral changes for the double and triple mutant retinochromes relative to the wildtype generally did not reflect additive changes of the single mutant effects. Together, these results indicate that amino acid substitutions in these spectral tuning sites may experience epistatic effects, the non-additive effect on protein function arising from identical amino acid substitutions in different genetic backgrounds. Intramolecular epistasis has been implicated in opsin evolution, likely resulting from trade-offs between selection maintaining protein stability and spectral tuning [51,52]. Selection on amino acid sites relating to properties unrelated to spectral tuning in retinochrome may therefore hinder our ability predict changes to absorbance from these three residues alone.

### Scallop retinochrome as a promising model for understanding opsin function

In this study we were able to identify and successfully mutate three amino acid sites that affect absorption maxima in scallop retinochrome. Our results demonstrate that retinochrome is easy to clone, mutate, and express in heterologous cell culture, making it as a promising model to examine the genotype-phenotype relationship in opsins. To make retinochrome a more powerful system, we need detailed atomic-level information to better model how all-*trans* retinal interacts with the protein and more extensive sampling across molluscan species to describe the natural variants and patterns of molecular evolution. This study and others [5,53,54] have shown that the amino acid sequence does not provide a reliable predictor of absorption maxima (λ_max_) trends. In fact, λ_max_ values are likely determined by changes in the topographical distribution and orientation of the amino acid side-chains surrounding the chromophore, making 3D structure of the photopigment essential. The application of Quantum Mechanics/Molecular Mechanics (QM/MM) models is a powerful tool for studying proteins beyond their structural characterizations [10,41,55,56]. However, the modeling of opsin proteins is particularly challenging as it not only requires an atomic-level representation to correctly describe the interaction between the chromophore and its environment, but also a correct description of its light-responsive electronic structure. Integrating experimental measurements of retinochrome mutants with model refinements could yield more accurate and consistent QM/MM models. With such tools, it will be possible to classify the effect of different noncovalent interactions, construct a topographical map of their distribution, and ultimately, develop hypotheses on how the λ_max_ values and reactivity properties of the photopigment respond to these interactions ([57]).

## Conclusions

We demonstrate the first example of expression, mutagenesis, and spectral analysis of retinochrome from scallop species. We showed that the absorbance peaks of scallop retinochromes fall within the blue-green range of the visible spectrum similar to *in vivo* measurements of cephalopod retinochrome. We propose methods to identify amino acid sites potentially responsible for spectral tuning of photopigments. Amino acids near sites of import, such as the Schiff base binding site and counterion, have long been recognized for their role in spectral tuning, but our results show that sites elsewhere in the protein may be responsible for tuning in the λ_max_. While we uncovered sites for spectral tuning, we rarely observed mirrored changes in λ_max_ for our reciprocal mutagenesis studies, suggesting that retinochrome function may be influenced by epistasis, thus complicating the predictability of genotype to phenotype relationships for this opsin. Finally, our findings highlight the effectiveness of scallop retinochrome as a model system for investigating the relationship between genotype and phenotype of photopigments.

## Materials and methods

### Retinochrome Cloning and Insertion in Vector

Previously assembled transcriptomes (Serb et al., *in prep*) were used to identify retinochrome sequences in *Argopecten irradians* and *Pecten maximus*. Based on those transcripts, UTR-specific primers were designed to amplify the complete coding regions from cDNA (Table 1). Scallop RNA was extracted from eye tissue using the RiboPure RNA extraction kit (Ambion) and converted to cDNA libraries. PCR was carried out with a reaction mixture equaling 50uL, containing, 5uL of 10x buffer, 1.5uL of 25mM MgCl, 4uL of 2.5mM dNTPs, 0.2uL Platinum Taq, 1uL of 10uM of forward and reverse primers (Table 1), and 1uL of 3uM template cDNA. The thermocycler protocol used was as follows, with variation in primer annealing temperatures: 95°C for 2min; 35 cycles of 95°C for 30s, primer temperature for 40s, and 72°C for 2min; and 72°C for 10 min. PCR products were size-screened using a 1% agarose gel electrophoresis, bands of expected size (923bp) were gel extracted (Qiagen Qiaquick Gel Extraction kit) and cloned using chemically competent *E. coli* cells following the manufacturer’s protocol (TOPO TA Cloning Kit with pCR2.1-TOPO). The identity of positive colonies from blue-white screening was confirmed by Sanger DNA sequencing using an ABI 3730 Capillary Electrophoresis Genetic Analyzer at the Iowa State University DNA Facility. The genes were then inserted into the expression vector p1D4-hrGFP II [59] to generate our working plasmids for retinochrome from *A. irradians* (Airr-RTC) and *P. maximus* (Pmax-RTC). These expression plasmids served as the templates for site-directed mutagenesis.

### Modeling and Site Identification

To identify amino acids lining the binding pocket which may be responsible for altering the λ_max_ of retinochrome, amino acid sequences and predicted 3D models were compared. Amino acid sequences of *A. irradians* retinochrome and *P. maximus* retinochrome were aligned using MAFFT v7.221 [60]. To identify sites hypothesized to cause changes in λ_max_ of retinochrome, amino acid sequences of *A. irradians* and *P. maximus* retinochrome were submitted to GPCR-I-TASSER [61] to create 3D models of each protein. The resulting models were then submitted to COACH [62,63], a meta-server used to predict the active interaction sites within protein-ligand interactions. COACH outputs a list of amino acid sites it has predicted to interact with the ligand when bound based on proximity of the amino acid to the bound ligand model. This list of predicted sites was compared to the alignment of Airr-RTC and Pmax-RTC, specifically to identify predicted interaction sites that are also not conserved sites between the two species, revealing one predicted interaction site which differed between species.

The second approach to identifying amino acids responsible for altering the retinochrome binding pocket environment was based on the role of possible polar interactions between amino acids and variation in the shape or electrostatic environment of the binding pocket plays a role in spectral tuning of the λ_max_ of opsins. 3D models from COACH were loaded into UCSF Chimera v1.4 [64], a visualization software for molecular analyses and model comparison. Using Chimera, predicted interaction sites were differentially highlighted based on whether the amino acids were conserved between *A. irradians* and *P. maximus* amino acid sequences. Non-conserved amino acids outside of or far from the binding pocket were disregarded, as they are less likely to affect the polar or shape of the binding pocket. The distances of the predicted interaction sites to the active side chains of non-conserved amino acids were then individually measured. Distances less than 3.5 angstroms (maximal hydrogen bond length) were searched for, revealing two sites as targets for site-directed mutagenesis.

### Site-directed Mutagenesis

To create fourteen mutants (all possible combinations of the three sites of interest explained previously as well as the reciprocal mutants), the Airr-RTC and Pmax-RTC expression plasmid served as the templates for the preceding mutagenesis experiments. DNA Polymerase PfuTurbo (Agilent, Santa Clara, CA) was used for all cloning experiments following the manufacturer’s instructions with the same thermocycler profile with varying annealing temperatures dependent on the specifications of each primer set (Supp. Table 1). Following PCR protocol, reaction was subjected to a 2.5-h digestion with DpnI (New England Biolabs, Ipswich, MA) at 37°C. Five microliters of the reaction was then used to transform TOP10 chemically competent *E. coli* cells (Thermo Fisher Scientific, Waltham, MA). Positive colonies containing the correct mutant sequence were confirmed by Sanger sequencing. Overlapping primers were developed to create mutant Airr-RTC and Pmax-RTC (Supp. Table 1). Single mutant primers were used in the creation of all mutants except for one double mutant plasmid. Due to the proximity of the selected sites, a set of primers including two mutation sites was used to guarantee mutation of both sites without removal of either. Mutant products were then used as templates for subsequent mutagenesis. Plasmids were amplified by incubating positively identified colonies in 1L liquid LB culture with 50 ug/mL kanamycin. Plasmids were purified using QIAGEN (Hilden, Germany) HiSpeed Plasmid Maxi Kit according to the manufacturer’s instructions with the product sequenced to confirm mutant identity.

The seven mutant proteins of Airr-RTC were successfully created, comprising all the combinations of mutations at the respective sites. Wild-type Airr-RTC was used as the backbone template for the single mutant proteins. Mutants are labeled with a delta indicating presence of a mutation followed the wild-type and resulting amino acid, respectively, flanking the site number: Airr-RTC-ΔM181L, Airr-RTC-ΔY188W, and Airr-RTC-ΔV193A. Double mutant proteins used the single mutant retinochromes as a template using the same primers used to create the single mutants; however, due to the proximity of sites 188 and 193, new overhang primers were designed to include the mutations at both sites (Supp. Table 1). This ensured the production of a double mutant without the chance of reverse mutation. The resulting mutants are: Airr-RTC-ΔM181L-ΔY188W, Airr-RTC-ΔM181L-ΔV193A, Airr-RTC-ΔY188W-ΔV193A. Finally, the production of the triple mutant was carried out using Airr-RTC-ΔY188W-ΔV193A as the template and the primer set for site 181: Airr-RTC- ΔM181L-ΔY188W-ΔV193A. Reciprocal mutants were made in the same manner using different primers (Supp. Table 1) to create the respective combinations of mutants with Pmax-RTC: Pmax-RTC-ΔL181M, Pmax-RTC-ΔW188Y, and Pmax-RTC-ΔA193V, Pmax-RTC-ΔL181M-ΔW188Y, Pmax-RTC-ΔL181M-ΔA193V, Pmax-RTC-ΔW188Y-ΔA193V, and Pmax-RTC- ΔL181M-ΔW188Y-ΔA193V.

### Cell Culture, Expression, and Pull-down

To express wild-type and mutant retinochrome proteins *in vitro*, 15 plates (Corning Falcon Standard Tissue Culture Dishes, 10cm, ref. 353003; Tewksbury, MA) of confluent HEK293T cells (ATCC, Manassas, VA) were transfected with 8 mg DNA and 20 mL 293fectin Transfecting Reagent (Thermo Fisher Scientific) per plate, according to the manufacturer’s instructions. Bovine rhodopsin was used as a system control using two plates of confluent HEK293T cells with equivalent amounts of DNA and 293fectin as described above. Plates were incubated for 24 hours before the minimum essential medium (MEM) was exchanged with new MEM containing 5 μmol all-*trans* retinal for retinochrome and 5 μmol of 11-*cis* retinal for bovine rhodopsin. Due to the addition of light sensitive retinal at this step, all subsequent culturing and experimentation was conducted in a dark-lab environment under dim red light. The plates were incubated for another 24 hours before cells were harvested by scraping the plates twice with 5 mL bufferA (3 mmol MgCl2, 140 mmol NaCl, 50 mmol HEPES pH 6.6, aprotinin [10 mg/mL], leupeptin [10 mg/mL]). All subsequent centrifugation and incubation steps were at 4°C or on ice. Cells were collected by pellet following centrifugation (10 min at 1620 relative centrifugal force [RCF]) and resuspended in 10 mL buffer A. Cells were washed two times in total following the same protocol.

After a second wash, cells were resuspended in 2 mL per plate of buffer A with 5 μmol all-*trans* retinal to regenerate the photopigment. The cell suspension was nutated for 1 hour at 4°C. The regenerated cells were pelleted by centrifugation for collection at 38360 RCF for 20 minutes and resuspended in solubilization buffer (buffer A plus 1% n-dodecyl b-D-maltoside and glycerol [20% w/v]) using 1 mL solubilization buffer per plate. The solubilized cells were nutated for 1 hour at 4C. After the hour nutation, the mixture was centrifuged for 20 min at 42,740 RCF. The supernatant was then added to a 100 mL slurry resin (1:1 v/v resin/resin buffer) composed of 1D4 antibody (University of British Columbia, Canada) conjugated to sepharose beads and nutated for 30 minutes. The resin was washed three times with 5 mL washing buffer (buffer A with 1% n-dodecyl b-D-maltoside and glycerol [20% w/v] without aprotinin and leupeptin), and the protein was eluted with 2 mL elution buffer (washing buffer with 40 mmol Rho1D4 peptide [TETSQVAPA]), adapted from Oprian et. al. (1987). To concentrate the protein sample, eluate was concentrated to ~300 uL using 4 Amicon Ultra 0.5 mL 10 kDa centrifugal filters (Millipore, Billerica, MA).

### Spectrophotometry

Ultraviolet-visible absorption spectra (250–750 nm) of purified proteins was measured at 15°C using a Hitachi U-3900 spectrophotometer (Chiyoda, Tokyo, Japan). Data analysis was performed on the mean value of five spectral measurements with the software UV Solutions v4.2 (Hitachi). To test the proteins for photoreactivity, “dark” absorbance was measured first for each protein, e.g., the naïve protein that has been incubated and regenerated with the all-*trans* retinal chromophore. Retinochrome proteins were tested independently with all-*trans* retinal because (1) retinochrome preferentially binds all-*trans* retinal [31] and (2) retinochrome forms a stable pigment only in the presence of all-*trans* retinal [37]. The maximum absorbance of the all-*trans* retinal when unbound to retinochrome apoprotein is 380 nm. Thus, any light-dependent isomerization converting free all-*trans* retinal to 11-*cis* retinal will be undetectable in the experimental system. Therefore, the most plausible explanation for any observed change in spectral absorbance is due to a conformational change of the retinal covalently bonded to the apoprotein.

For the “light” spectra, extracted proteins were bleached at different wavelengths according to λ_max_ identified from the “dark” spectra and then the absorbance was measured. Extracted proteins were first exposed to light at ∼474 nm using two blue LEDs (MR16-B24-15-DI; http://superbrightleds.com) simultaneously irradiating both transparent sides of the cuvette (Hellma Analytics 104002B-10-40; Müllheim, Germany) for 3 minutes and the absorbance was recorded. A final exposure to white light was performed, followed by an absorbance measurement. Mean spectra was plotted using R scripts (R Core Team, 2017) with interpolation of the data points being performed with the R function smooth.spline. The differential spectra were calculated from two adjacent light treatments (i.e., the absorbance spectrum before exposure to light [dark spectra] was subtracted from spectra after the exposure to blue light [blue spectra]). This approach minimized the likelihood of observed differences in spectrum being a result of unrelated factors such as handling of the cuvette between light exposures and absorbance measurements or degradation of the protein sample other than light treatments themselves. These methods were carried out for the wild-type proteins of *A. irradians* and *P. maximus* and then the seven mutant *A. irradians* proteins and seven mutant *P. maximus* proteins.

## Acknowledgments

We thank Rosalie Crouch (Storm Eye Institute, Medical University of South Carolina) and the National Eye Institute for supplying 11-*cis*-retinal, Belinda Chang for providing the expression vector p1D4-hrGFP II, and Julia Sigwart for scallop specimens. We are indebted to Davide Faggionato, who developed the heterologous expression method for scallop opsins. We are grateful to the Serb lab for comments on earlier drafts, in particular Jorge Audino. This study is part of GDS’s PhD thesis through the Graduate Program in Interdepartmental Genetics and Genomics (Iowa State University).

## Author Contributions

GDS and JMS are responsible for conceptualization and methodology of this study. GDS and KEM conducted the investigation and produced mutant genes and cloning of expression plasmids. GDS and KEM carried out all protein expressions, protein isolation and spectral analyses. GDS produced all figures and wrote the original manuscript with review and edits in collaboration with KEM and JMS. JMS was responsible for funding acquisition.

## References

1. Shichida Y, Matsuyama T. Evolution of opsins and phototransduction. Philos Trans R Soc Lond B Biol Sci. 2009;364: 2881–2895. doi:10.1098/rstb.2009.0051

2. Ueyama H, Kuwayama S, Imai H, Tanabe S, Oda S, Nishida Y, et al. Novel missense mutations in red/green opsin genes in congenital color-vision deficiencies. Biochem Biophys Res Commun. 2002;294: 205–209. doi:10.1016/S0006-291X(02)00458-8

3. Beppu Y, Kakitani T. Theoretical study of color control mechanism in retinal proteins. 1994;59: 660–669.

4. Beppu Y. Theoretical Study of color control mechanism in retinal proteins: orientational effects of aromatic amino acid residues upon opsin shift. 1997. pp. 3303–3309.

5. Nathans J. Determinants of visual pigment absorbance: Identification of the retinylidene schiff’s base counterion in bovine rhodopsin. Biochemistry. 1990;29: 9746–9752. doi:10.1021/bi00493a034

6. Oprian DD, Pelletier SL, Asenjo AB, Lee N. Design, chemical synthesis, and expression of genes for the three human color vision pigments. Biochemistry. 1991;30: 11367–11372. doi:10.1021/bi00112a002

7. Yokoyama S. Phylogenetic analysis and experimental approaches to study color vision in vertebrates. Methods Enzymol. 2000;315: 312–325. doi:10.1016/s0076-6879(00)15851-3

8. Yokoyama S. Molecular evolution of vertebrate visual pigments. Progress in Retinal and Eye Research. 2000. doi:10.1016/S1350-9462(00)00002-1

9. Van Hazel I, Sabouhanian A, Day L, Endler JA, Chang BS. Functional characterization of spectral tuning mechanisms in the great bowerbird short-wavelength sensitive visual pigment (SWS1), and the origins of UV/violet vision in passerines and parrots. BMC Evol Biol. 2013;13. doi:10.1186/1471-2148-13-250

10. Melaccio F, Ferré N, Olivucci M. Quantum chemical modeling of rhodopsin mutants displaying switchable colors. Phys Chem Chem Phys. 2012;14: 12485–12495. doi:10.1039/c2cp40940b

11. Ferré N, Olivucci M. Probing the rhodopsin cavity with reduced retinal models at the CASPT2/CASSCF/AMBER level of theory. J Am Chem Soc. 2003;125: 6868–6869. doi:10.1021/ja035087d

12. González-Luque R, Garavelli M, Bernardi F, Merchán M, Robb MA, Olivucci M. Computational evidence in favor of a two-state, two-mode model of the retinal chromophore photoisomerization. Proc Natl Acad Sci U S A. 2000;97: 9379–9384. doi:10.1073/pnas.97.17.9379

13. Irving CS, Byers GW, Leermakers PA. Spectroscopic model for the visual pigments: Influence of microenvironmental polarizability. Biochemistry. 1970;9: 858–864. doi:10.1021/bi00806a020

14. Lin SW, Kochendoerfer GG, Carroll KS, Wang D, Mathies RA, Sakmar TP. Mechanisms of spectral tuning in blue cone visual pigments: Visible and raman spectroscopy of blue-shifted rhodopsin mutants. J Biol Chem. 1998;273: 24583–24591. doi:10.1074/jbc.273.38.24583

15. Merbs SL, Nathans J. Role of hydroxyl-bearing amino acids in differentially tuning the absorption spectra of the human red and green cone pigments. Photochem Photobiol. 1993;58: 706–710. doi:10.1111/j.1751-1097.1993.tb04956.x

16. Hauser FE, van Hazel I, Chang BSW. Spectral tuning in vertebrate short wavelength-sensitive 1 (SWS1) visual pigments: Can wavelength sensitivity be inferred from sequence data? J Exp Zool Part B Mol Dev Evol. 2014;3228: 529–539. doi:10.1002/jez.b.22576

17. Hunt DM, Carvalho LS, Cowing JA, Davies WL. Evolution and spectral tuning of visual pigments in birds and mammals. Philos Trans R Soc B Biol Sci. 2009;364: 2941–2955. doi:10.1098/rstb.2009.0044

18. Hope AJ, Partridge JC, Dulai KS, Hunt DM. Mechanisms of wavelength tuning in the rod opsins of deep-sea fishes. Proc R Soc B Biol Sci. 1997;264: 155–163. doi:10.1098/rspb.1997.0023

19. Chan T, Lee M, Sakmar TP. Introduction of hydroxyl-bearing amino acids causes bathochromic spectral shifts in rhodopsin. Amino acid substitutions responsible for red-green color pigment spectral tuning. J Biol Chem. 1992;267: 9478–9480.

20. Asenjo AB, Rim J, Oprian DD. Molecular determinants of human red/green color discrimination. Neuron. 1994;12: 1131–1138. doi:10.1016/0896-6273(94)90320-4

21. Neitz M, Neitz J, Jacobs GH. Spectral tuning of pigments underlying red-green color vision. Science (80-). 1991;252: 971 LP – 974. doi:10.1126/science.1903559

22. Yokoyama S, Bernhard Radlwimmer F. The “five-sites” rule and the evolution of red and green color vision in mammals. Mol Biol Evol. 1998;15: 560–567. doi:10.1093/oxfordjournals.molbev.a025956

23. Yokoyama S, Bernhard Radlwimmer F. The molecular genetics of red and green color vision in mammals. Genetics. 1999;153: 919–932.

24. Hunt DM, Carvalho S, Cowing JA, Parry JWL, Wilkie SE, Davies WL, et al. Spectral tuning of shortwave-sensitive visual pigments in vertebrates †. Photochem Photobiol. 2007;83: 303–310. doi:10.1562/2006-06-27-IR-952

25. Yokoyama S. Evolution of dim-light and color vision pigments. Annu Rev Genomics Hum Genet. 2008;9: 259–282. doi:10.1146/annurev.genom.9.081307.164228

26. Wakakuwa M, Terakita A, Koyanagi M, Stavenga DG, Shichida Y, Arikawa K. Evolution and mechanism of spectral tuning of blue-absorbing visual pigments in butterflies. PLoS One. 2010;5: 1–8. doi:10.1371/journal.pone.0015015

27. Saito T, Koyanagi M, Sugihara T, Nagata T, Arikawa K, Terakita A. Spectral tuning mediated by helix III in butterfly long wavelength-sensitive visual opsins revealed by heterologous action spectroscopy. Zool Lett. 2019;5: 1–11. doi:10.1186/s40851-019-0150-2

28. Terakita A, Tsukamoto H, Koyanagi M, Sugahara M, Yamashita T, Shichida Y. Expression and comparative characterization of Gq-coupled invertebrate visual pigments and melanopsin. J Neurochem. 2008;105: 883–890. doi:10.1111/j.1471-4159.2007.05184.x

29. Varma N, Mutt E, Mühle J, Panneels V, Terakita A, Deupi X, et al. Crystal structure of jumping spider rhodopsin-1 as a light sensitive GPCR. Proc Natl Acad Sci U S A. 2019;116: 14547–14556. doi:10.1073/pnas.1902192116

30. Shimamura T, Hiraki K, Takahashi N, Hori T, Ago H, Masuda K, et al. Crystal structure of squid rhodopsin with intracellularly extended cytoplasmic region. J Biol Chem. 2008;283: 17753–17756. doi:10.1074/jbc.C800040200

31. Hara T, Hara R. Rhodopsin and Retinochrome in the Squid Retina. Nature. 1967;214.

32. Hara T, Hara R, Hara NI, Nishimura M, Terakita A, Ozaki K. The rhodopsin-retinochrome system in the squid visual cell. Colloque INSERM; Structures et fonctions des retino proteines Colloquium; Structures and functions of retinal proteins. 1992. pp. 255–258.

33. Hara T, Hara R. Cephalopod Retinochrome. Photochem Vis. 1972;7: 720–746. doi:10.1016/S0076-6879(82)81031-8

34. Ozaki K, Hara R, Hara T, Kakitani T. Squid retinochrome. Configurational changes of the retinal chromophore. Biophys J. 1983;44: 127–137. doi:10.1016/S0006-3495(83)84285-4

35. Hara T, Hara R. Regeneration of Squid Retinochrome. Nature. 1968;219: 450–454. doi:10.1038/219450a0

36. Hara T, Hara R, Takeuchi J. Rhodopsin and Retinochrome in the Octopus Retina. Nature. 1967;214: 572–573. doi:10.1038/214572a0

37. Faggionato D, Serb JM. Strategy to identify and test putative light-sensitive non-opsin G-protein-coupled receptors: A case study. Biol Bull. 2017;233: 70–82. doi:10.1086/694842

38. Ramirez MD, Pairett AN, Pankey MS, Serb JM, Speiser DI, Swafford AJ, et al. The last common ancestor of most bilaterian animals possessed at least nine opsins. Genome Biol Evol. 2016;8: 3640–3652. doi:10.1093/gbe/evw248

39. Terakita A, Yamashita T, Shichida Y. Highly conserved glutamic acid in the extracellular IV-V loop in rhodopsins acts as the counterion in retinochrome, a member of the rhodopsin family. Proc Natl Acad Sci U S A. 2000;97: 14263–14267. doi:10.1073/pnas.260349597

40. Sperling L, Hubbard R. Squid retinochrome. J Gen Physiol. 1975;65: 235–251. doi:10.1085/jgp.65.2.235

41. Hirayama J, Imamoto Y, Shichida Y, Yoshizawa T, Asato AE, Liu RSH, et al. Shape of the chromophore binding site in pharaonis phoborhodopsin from a study using retinal analogs. Photochem Photobiol. 1994;60: 388–393. doi:10.1111/j.1751-1097.1994.tb05121.x

42. Rinaldi S, Melaccio F, Gozem S, Fanelli F, Olivucci M. Comparison of the isomerization mechanisms of human melanopsin and invertebrate and vertebrate rhodopsins. Proc Natl Acad Sci U S A. 2014;111: 1714–1719. doi:10.1073/pnas.1309508111

43. Ozaki K, Terakita A, Hara R, Hara T. Rhodopsin and retinochrome in the retina of a marine gastropod, Conomulex luhuanus. Vision Res. 1986;26: 691–705. doi:10.1016/0042-6989(86)90083-0

44. Shimono K, Ikeura Y, Sudo Y, Iwamoto M, Kamo N. Environment around the chromophore in *Pharaonis* phoborhodopsin: Mutation analysis of the retinal binding site. Biochim Biophys Acta - Biomembr. 2001;1515: 92–100. doi:10.1016/S0005-2736(01)00394-7

45. Nakayama TA, Khorana HG. Mapping of the amino acids in membrane-embedded helices that interact with the retinal chromophore in bovine rhodopsin. J Biol Chem. 1991;266: 4269–4275.

46. Sekharan S, Katayama K, Kandori H, Morokuma K. Color vision: “OH-site” rule for seeing red and green. J Am Chem Soc. 2012;134: 10706–10712. doi:10.1021/ja304820p

47. Collette F, Renger T, Müh F, Schmidt Am Busch M. Red/green color tuning of visual rhodopsins: Electrostatic theory provides a quantitative explanation. J Phys Chem B. 2018;122: 4828–4837. doi:10.1021/acs.jpcb.8b02702

48. Greenhalgh DA, Farrens DL, Subramaniam S, Khorana HG. Hydrophobic amino acids in the retinal-binding pocket of bacteriorhodopsin. J Biol Chem. 1993;268: 20305–20311.

49. Surridge AK, Osorio D, Mundy NI. Evolution and selection of trichromatic vision in primates. Trends Ecol Evol. 2003;18: 198–205.

50. Neitz M, Neitz J, Jacobs GH. Spectral tuning of pigments underlying red-green color vision Author (s): Maureen Neitz , Jay Neitz and Gerald H . Jacobs Published by : American Association for the Advancement of Science Stable URL : http://www.jstor.org/stable/2875359 Accessed : 07-0. 2016;252: 971–974.

51. Dungan SZ, Chang BSW. Epistatic interactions influence terrestrial–marine functional shifts in cetacean rhodopsin. Proc R Soc B Biol Sci. 2017;284. doi:10.1098/rspb.2016.2743

52. Castiglione GM, Chang BSW. Functional trade-offs and environmental variation shaped ancient trajectories in the evolution of dim-light vision. Elife. 2018;7: 1–30. doi:10.7554/eLife.35957

53. Yokoyama S, Xing J, Liu Y, Faggionato D, Altun A, Starmer WT. Epistatic adaptive evolution of human color vision. PLoS Genet. 2014;10. doi:10.1371/journal.pgen.1004884

54. Yokoyama S, Altun A, Jia H, Yang H, Koyama T, Faggionato D, et al. Adaptive evolutionary paths from UV reception to sensing violet light by epistatic interactions. Sci Adv. 2015;1. doi:10.1126/sciadv.1500162

55. Ferre N, Olivucci M. Energy storage of rhodopsin resolved at the multiconfigurational perturbation theory level. 2004;101: 17908–17913.

56. Coto PB, Strambi A, Ferre N, Olivucci M. The color of rhodopsins at the ab initio multiconfigurational perturbation theory resolution. Proc Natl Acad Sci U S A. 2006;103: 17154–17159. doi:10.1073/pnas.0604048103

57. Gozem S, Melaccio F, Luk HL, Rinaldi S, Olivucci M. Learning from photobiology how to design molecular devices using a computer. Chem Soc Rev. 2014;43: 4019–4036. doi:10.1039/c4cs00037d

58. Serb JM, Faggionato D, Smedley GD, Seetharam A, Severin AJ, Pairett AN. Expression and spectral analysis of twelve opsins reveals astonishing photochemical diversity in the common bay scallop Argopecten irradians (Mollusca: Bivalvia).

59. Morrow JM, Chang BSW. The p1D4-hrGFP II expression vector: a tool for expressing and purifying visual pigments and other G protein-coupled receptors. Plasmid. 2010;64: 162–169. doi:10.1016/j.plasmid.2010.07.002

60. Katoh K, Standley DM. MAFFT multiple sequence alignment software version 7: Improvements in performance and usability. Mol Biol Evol. 2013;30: 772–780. doi:10.1093/molbev/mst010

61. Zhang J, Yang J, Jang R, Zhang Y. GPCR-I-TASSER: A hybrid approach to G Protein-Coupled Receptor structure modeling and the application to the human genome. Structure. 2015;23: 1538–1549. doi:10.1016/j.str.2015.06.007

62. Yang J, Roy A, Zhang Y. Protein-ligand binding site recognition using complementary binding-specific substructure comparison and sequence profile alignment. Bioinformatics. 2013;29: 2588–2595. doi:10.1093/bioinformatics/btt447

63. Yang J, Roy A, Zhang Y. BioLiP: A semi-manually curated database for biologically relevant ligand-protein interactions. Nucleic Acids Res. 2013;41: 1096–1103. doi:10.1093/nar/gks966

64. Pettersen EF, Goddard TD, Huang CC, Couch GS, Greenblatt DM, Meng EC, et al. UCSF Chimera - A visualization system for exploratory research and analysis. J Comput Chem. 2004;25: 1605–1612. doi:10.1002/jcc.20084

65. Oprian DD, Molday RS, Kaufman RJ, Khorana HG. Expression of a synthetic bovine rhodopsin gene in monkey kidney cells. Proc Natl Acad Sci U S A. 1987;84: 8874–8. doi:10.1073/pnas.84.24.8874

